# “Decoding Chromosome Radiosensitivity in G_0_-Phase PBMCs: Genomic Determinants, Age, Sex effects and Implications in Radiation Dose Assessment”

**DOI:** 10.64898/2026.05.31.728688

**Authors:** Usha Yadav, Utkarsha S Mungse, Nagesh N Bhat, Arshad Khan, Balvinder K Sapra

## Abstract

Low-dose ionizing radiation from natural and anthropogenic sources is typically not of significant concern under normal conditions. However, in case of radiological incidents, it becomes an important environmental hazard, raising concern for public health and necessitating reliable biological indicators of exposure.

Chromosomal aberrations are most reliable markers of radiation exposure and are more or less universally assessed in metaphases. Recently, rapid assessment of aberrations in G₀-phase lymphocytes using G₀-PCC-FISH has gained attention and is recognized as the most suitable method for dose assessment in cases of high-dose or partial-body exposure.

This is first study in G₀-phase at such scale to establish baseline and radiation induced aberrations in peripheral blood mononuclear cells (PBMCs) from 24 healthy human donors (12 males, 12 females, 21-60 Y). Using whole chromosome painting (WCP) of Chromosomes 1, 2 & 4 and scoring across >8500 cells post 0, 2, and 4 Gy γ-radiation exposure, we quantified 5586 aberrations and investigated their relationship with underlying genomic features.

**Our findings include:** No significant effect of age or sex on chromosomal radiosensitivity, supporting the robustness of pooled biodosimetric calibration curves for dose assessment within the studied age range of 21–60 years.

Chromosome-specific radiosensitivity does not appear to be solely dependent on chromosome size and shows a potential association with gene density and total transcript length.

From public health point of view the present data provides reference values for interphase chromosomal damage as well as radiation induced reference values for two important dose points across age groups and sexes. This approach enhances emergency preparedness for radiological events by enabling rapid biodosimetry, especially critical when metaphase cells are unavailable as in cases of accidental high-dose or partial exposures.

Graphical Abstract:
A 24-donor study to assess baseline and radiation response age and sex. Additionally substantial data analysed to assess radiosensitivity of chromosomes and its genomic determinants.

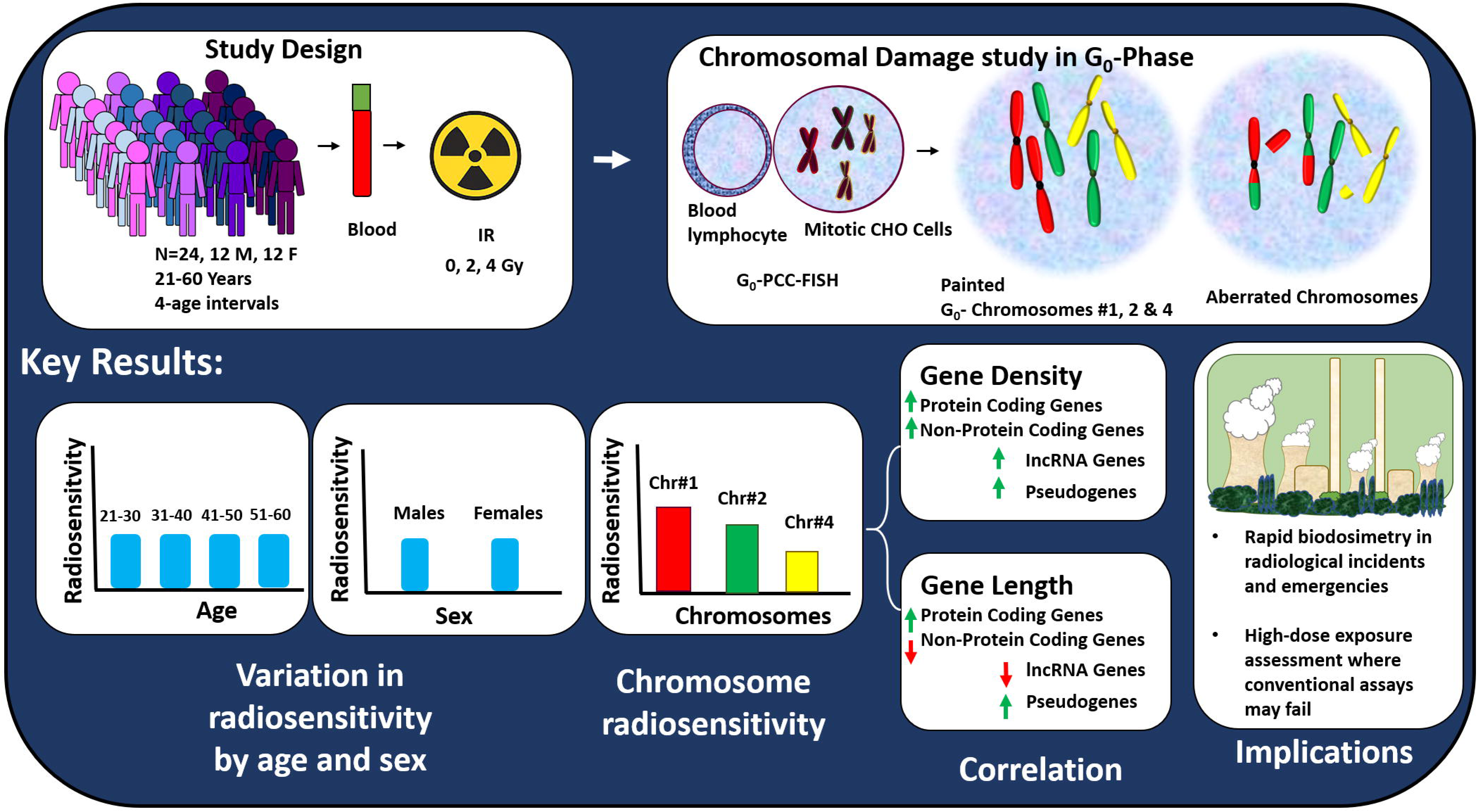

## 1. Introduction

Ionizing radiation is an intrinsic component of the environment, contributing to continuous low-dose exposure from natural background sources as well as anthropogenic activities including medical procedures, industrial applications and nuclear power generation. However, accidental events such as Chernobyl, Fukushima and the Goiania incident can lead to elevated environmental radiation levels with significant ecological and public health consequences. In addition, populations residing in high natural background radiation areas and as well as individuals exposed through medical or occupational settings require careful evaluation to understand potential biological effects. These scenarios emphasize the importance of developing sensitive and reliable methods and biological indicators to assess radiation-induced damage and to support environmental monitoring of exposed populations.

Genomic integrity is fundamental to human health and prevalence of small to large scale mutations or aberrations is indicator of genomic instability and a central driver of carcinogenesis, aging, degenerative diseases and genetic abnormalities (Bonassi et al., 2000, 2008; Jacobs et al., 2012; H. Wang et al., 2017; S. S. Wang et al., 2008).

It is known that radiation exposure can induce DNA damage and aberrations in a dose dependent manner which serve as standard biological indicators of radiation exposure. Detecting and quantifying these aberrations in human cells offers insights into both the absorbed radiation dose (biodosimetry) in case of suspected radiation over-exposure to public or radiation workers and individual susceptibility to radiation. Conventionally, the standard approach to evaluating chromosomal aberrations has involved culturing peripheral blood lymphocytes for metaphase-based analysis (IAEA, 2011). While effective, this method is time-consuming (requiring 48–72 hours), and dependent on the ability of lymphocytes to reach metaphase. While powerful, metaphase-based assays do not capture early DNA damage events or repair kinetics. In recent years in genotoxicity assessment, premature chromosome condensation (PCC) combined with fluorescence in situ hybridization (FISH) has emerged as a powerful tool. It enables rapid analysis of chromosomal damage in G_0_-Phase cells, without the need for 48-hour culture. This approach allows for a more direct and potentially more accurate assessment of baseline DNA damage and radiosensitivity in the human population. It also enables for analysis of DNA repair kinetics, helping to identify individuals with defective repair pathways and higher radiation sensitivity (Hatzi et al., 2006; Karachristou et al., 2015; Pantelias & Terzoudi, 2018). Moreover, this technique can be applied to samples where metaphases are unavailable, such as high dose radiation exposure, or immune compromised patients. The technique has been found to have potential advantages over metaphase-based biodosimetry for accidental high-dose and partial body radiation exposure cases (Darroudi et al., 1998; Durante et al., 1998; Fomina, 2000; Fomina et al., 2001; Rao & Natarajan, 2001; Ryan et al., 2019; Yadav et al., 2021, 2024).

It is crucial to establish baselines of the aberrations among healthy individuals and possible confounding factors such as age & sex before applying the technique. The influence of age and sex at the cytogenetic level has been assessed in stimulated lymphocytes showing a trend for higher micronucleus frequency with increasing age. Some report a slight increase in translocation frequency with age whereas, there is lesser evidence of increase in sister chromatid exchanges and inconsistent data for structural chromosomal type aberrations with respect to age (Bolognesi et al., 1997; Pressl et al., 1999; Rastkhah et al., 2016; Santovito et al., 2016; Sigurdson et al., 2008; I. Vorobtsova et al., 2001; I. E. Vorobtsova et al., 2000; Whitehouse et al., 2005). Most of the available literature focuses on metaphase-based analysis, leaving a significant gap in our understanding of how these variables influence chromosomal aberrations in interphase cells. Moreover, while preliminary studies have suggested that radiation induced aberrations may be marginally higher in females using interphase assays, comprehensive investigations in large and diverse cohorts are lacking (Yadav et al., 2020).

Further, it is essential to establish radiosensitivities of the chromosomes among individuals across age & sex and understand genomic determinants of the chromosomal radiosensitvity. There is limited and inconsistent data on the radiosensitivity of different chromosomes. Most previous studies are carried out at metaphase and restricted to 1-3 donors hence, limited aberration data is available. Deviation from DNA-proportional distribution of radiation induced chromosome aberrations has been indicated in a few studies (Barquinero et al., 1998; Knehr et al., 1994, 1996; Luomahaara et al., 1999; Sommer et al., 2005). Biological basis of differential sensitivities is not well understood. To assess radiation effects more reliably, chromosomes 1, 2, and 4 are commonly targeted in 3-color FISH protocols. These three largest human chromosomes offer substantial genome coverage and yield statistically meaningful aberration data within a practical scoring time of less than 24 hours, making them well-suited for both research and potential clinical applications.

The present study aims to fill this gap by conducting a systematic evaluation of chromosome-specific aberrations in G_0_-phase PBMCs using G_0_-PCC-FISH. A thorough analysis has been carried out to assess the differential radiosensitivity of Chr1, 2 & 4. We have examined the correlation of chromosomal sensitivities with their genomic content, gene densities, different genes types to address biological basis of the differential radiation effect. The influence of dose, age and sex were assessed on individual chromosome sensitivities. Further, age- and sex-related variability in baseline as well as in the radiation response was assessed.

## 2. Materials & Methods

### 2.1. Materials

For, Fluorescence in situ hybridization (FISH), a cocktail of fluorescent probes against Chromosomes 1, 2 & 4 was obtained from Metasystems. Culture media; Dulbecco’s Modified Eagle Medium (DMEM) and Roswell Park Memorial Institute Medium (RPMI-1640), as well as Fetal Bovine Serum (FBS), Trypsin were procured from Gibco. For cell fusion, 50% w/v, Polyethylene Glycol (PEG) with MW 1450 was obtained from Sigma-Aldrich. Density Gradient media; Histopaque-1077, Colcemid powder were also purchased from Sigma. Glacial acetic acid, Methanol were purchased from Sisco Research Laboratories Pvt. Ltd., and Potassium Chloride was obtained from Thomas Baker.

### 2.2. Biological resource

Blood samples were collected from healthy consented donors, including both males and females, across four age groups: 21-30 years, 31-40 years, 41-50 years and 51-60 years. In total, 24 donors were included in the study, with 12 biological females and 12 biological males. Each age group included six volunteers, with three females and three males. Individuals with a history of smoking, blood transfusion, or any co-morbidity were excluded.

### 2.3. Isolation of PBMCs

Lymphocytes were separated using density gradient method. For this, diluted blood samples were layered on Ficoll density gradient media. The blood was diluted 1:1 with RPMI-1640 medium, and the ratio of diluted blood to Ficoll was also 1:1. The two layers remained unmixed and centrifuged at 400 x g in fixed bucket rotor at 25 ^◦^C. The buffy coat layer was isolated and washed with 10 ml RPMI supplement with 10% FBS. Further, the cells were resuspended in RPMI and 10% FBS. The cells were counted and irradiated with Co-60 gamma radiation and incubated for 24 h at 37 ^◦^C with 5% CO_2_.

### 2.4. Exposure to Co-60 gamma radiation

Gamma exposure was carried out at 1Gy/min dose rate to deliver 2 Gy and 5 Gy doses using Bhabhatron facility at Bhabha Atomic Research Centre. The exposure duration and transport of the sample to incubator was ∼15-min. The source was calibrated and measurements were traceable to primary standards.

### 2.5. Premature chromosome condensation(G_0_-PCC)

To study chromosomal aberrations at resting phase itself, chromosomes were artificially condensed using mitotic cell fusion induced G_0_-PCC method. Previously reported protocol for mitotic cell collection and cryopreservations and PCC was followed (Yadav et al., 2021). For this, mitotic cells were enriched from cultured Chinese hamster ovary cell line. Growing cells were arrested at metaphase using Colcemid 0.2 μg/ml and floating cells were harvested ∼4 h post exposure. The cells were cryopreserved in liquid nitrogen until use. On the day of PCC mitotic cells were recovered, mixed with PBMCs in 1:5 ratio and centrifuged. The cell pellet was treated with PEG for 2 min at room temperature. Cell suspension with PEG was diluted with 4 ml RPMI and cells were again centrifuged. The pellet was resuspended in 0.5 ml RPMI supplemented with 10% FBS, 1 μg/ml Colcemid. The suspension was incubated for 2 h at 37^◦^C with 5% CO_2_ for the chromosome condensation among fused mitotic-PBMCs. After 2 h the suspension was treated with 8 ml, hypotonic solution of 0.45% KCl followed by fixation in 3:1 Methanol: Acetic acid. Chromosomal spreads were then prepared onto clean, chilled, moist microscopic glass slides.

### 2.6. Fluorescent in-situ Hybridization

Chromosome specific painting was carried out using cocktail of whole chromosome paints for Chr1, 2 & 4 from Metasystems and assay was performed as per manufacturer’s protocol. The slides were covered with 10 μl probe, spread using coverglass and sides were sealed with rubber cement. The sealed slide was heated for denaturation at 75 ^◦^C for 2 min followed by overnight incubation at 37^◦^C in dark, in a humid chamber. Next day, coverglass and all traces of rubber cement were removed carefully. Slide were washed in 0.4X SSC (pH 7.0) at 72°C (±1°C) for 2 min. Followed by another wash in 2X SSC, 0.05% Tween-20 (pH 7.0) at room temperature for 30 seconds. Slides were then rinsed in distilled water and air dried. Counterstaining of all the chromosomes was carried out using 20 μl DAPI with antifade. The slides were then observed under microscope.

### 2.7. Imaging and Scoring

Imaging and scoring of the cells were carried out using Axio-imager Z2 microscope, equipped with Metafer-5, ISIS modules for metaphase capture and FISH analysis. For each individual ∼100 PCC spreads per dose point were scored manually on ISIS software. Two fluorescence signals with similar sizes for the three painted chromosomes were counted as normal. The excess fluorescent signals were counted as aberration and its frequency per cell was calculated for each individual. Frequency of aberrations were counted separately for the three chromosomes for assessing chromosome sensitivities. Each excess fluorescent signal counted as aberration was further annotated by their types i.e., fragments, translocations and insertions. Translocations were further classified as either one-way or two-way. We analysed individual contribution of these aberrations to the overall yield. Representative images of the differentially painted chromosomes in an unaberrated cell and different aberrations among the aberrated PCC spreads are displayed in Figure 1. Scoring was carried out by two trained individuals and was performed in a non-blinded manner.

**Figure 1.**
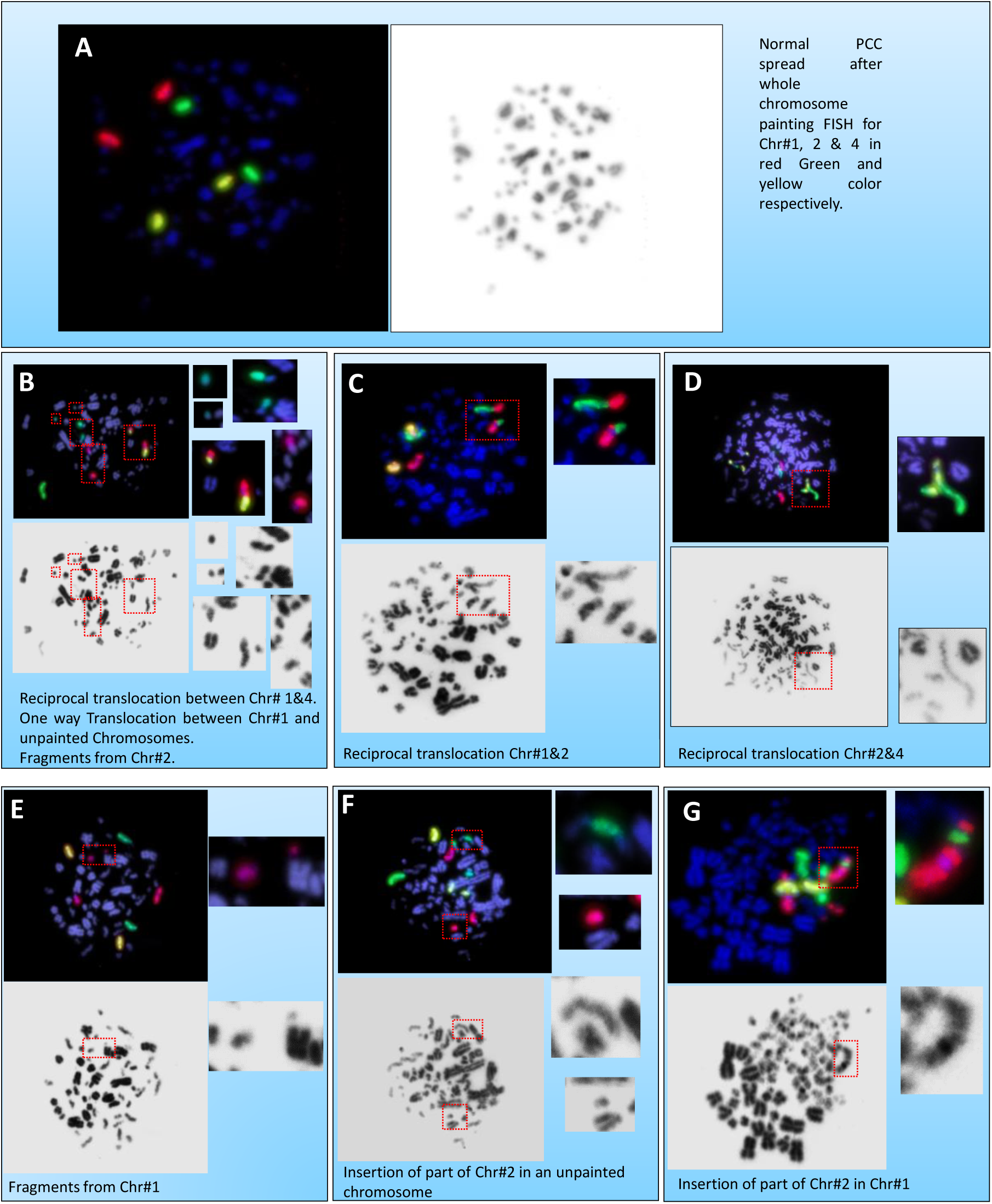
Representative images after G_0_-PCC-FISH in human PBMCs (A) A normal PCC spread showing two equal sized signals for the three painted chromosomes in red (Chr1), green (Chr2) and yellow (Chr4). (B-G) Annotation of different aberration types involving the painted chromosome.

### 2.8. Genomic Data

Genomic annotations for chromosomes 1, 2, and 4 were extracted separately from, NCBI,GRCh38.p14 database and processed using identical pipelines (accessed on 22 July 2025) (NCBI, 2025). Annotated genes were selected and details of transcripts were downloaded. All annotated transcripts, including both curated and predicted RefSeq entries were included. Multiple transcript isoforms per gene were included in the counts as well as length. Overlapping annotations were not explicitly filtered and were retained in the dataset (Supplementary data). Later data at gene level was also extracted by filtering duplicate entries per Gene IDs (Supplementary data).

### 2.9. Statistical Analysis

Poisson statistics were followed for calculating mean, standard error and significance testing for individual donor’s aberration counts and aberrations pooled from 24 donors (IAEA, 2011). Then, we computed mean aberration per donor and took the mean across 24 donors. For this mean of means, normal distribution statistics was followed, statistical significance was evaluated using the Student’s t-test.

## 3. Results & Discussion

### 3.1. Differential Chromosome Sensitivity towards Radiation Induced Aberrations: Chr1 is largest contributor of aberration at all the doses

Whole chromosome painting in differential colors of three different chromosomes allowed us to quantify the radiation sensitivity of the painted chromosomes separately. To achieve this, aberration data from over 8500 scored cells among 72-samples resulting from 0 Gy, 2 Gy and 4 Gy exposures for each of the 24 donors were analyzed. A total of 5586 aberrations were obtained. In overall results 40% of aberrations were associated with Chr1, 35% with Chr2 and 25 % with Chr4. We performed two sets of analysis, in one we pooled the aberrations from all the donors for the three chromosomes for each dose (Table 1). We calculated the aberration frequency per 100 cells and the percentage contribution of the three chromosomes to total aberrations Figure 2A & B. In the second analysis, mean frequency of aberrations were calculated for each donor for the three chromosome and three doses. A mean of the 24-individual means was then derived and plotted in Figure 2C. A heat map was generated to demonstrate individual variability in the radiosensitivity data Figure 2D. It can be observed that both pooled means and mean of means infer similar outcome. Both datasets demonstrate that the highest yield of aberrations is associated with Chr1 at all the doses while the lowest yields are observed for the Chr4.

**Figure 2.**
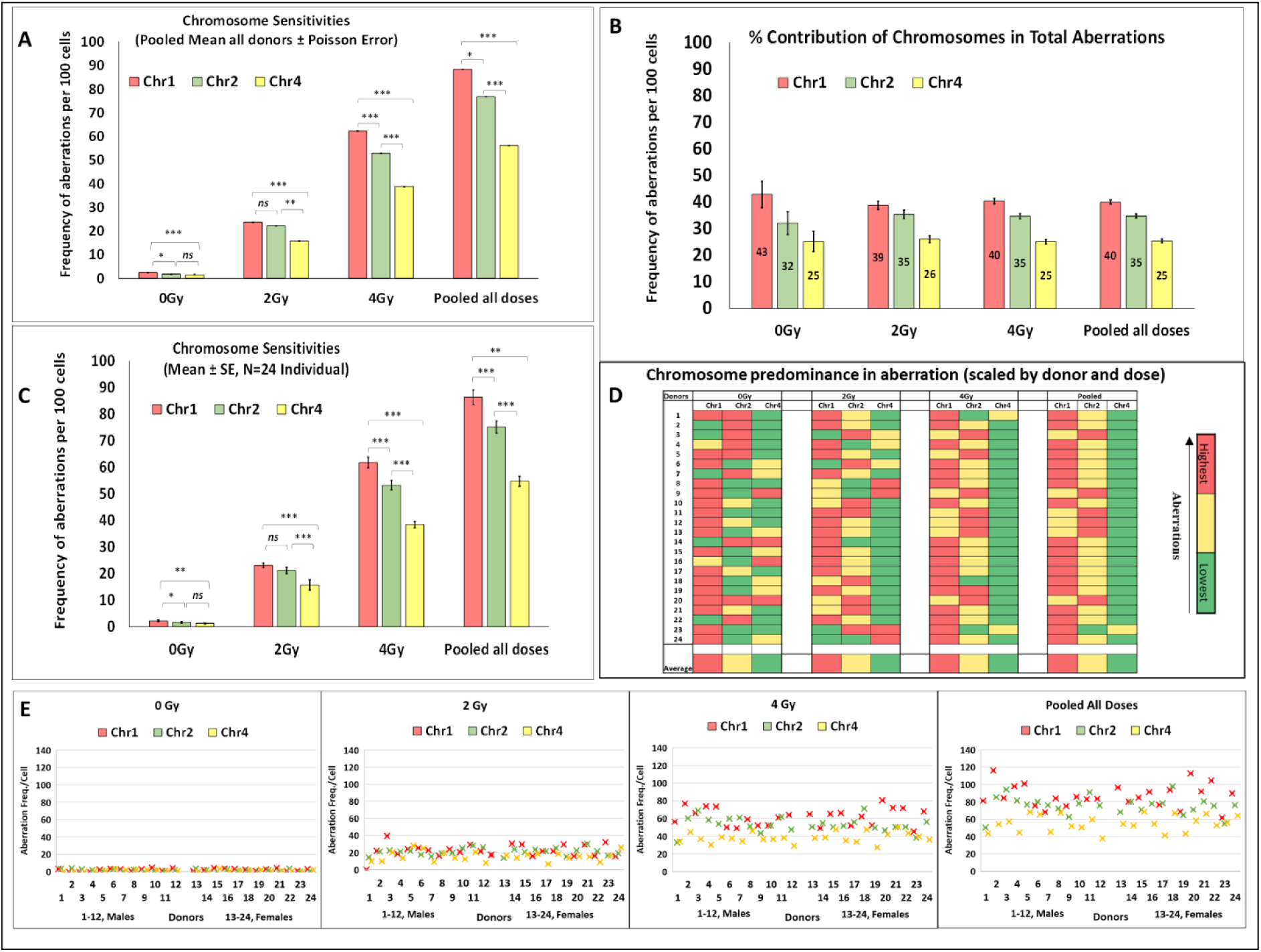
Chromosomal aberration frequency from Chr 1, 2, and 4 assessed using G₀-PCC-FISH based quantification across 3-radiation doses. Pooled all doses term refers to ‘sum of aberrations per 100 from all the doses’. A. Pooled chromosomal aberration data from 24 donors across varying radiation doses. B. % Contribution of aberrations by Chr1, 2, and 4 at different doses, based on pooled data from the 24 donors. C. Mean values of individual donor-level data (n = 24), showing consistent trends with the pooled dataset, confirming reproducibility. D. Heatmap showing relative aberration frequency (scaled by donor and dose separately) illustrating chromosomal predominance in the aberration. E. Scatter Plot of the 24-donors’ chromosomal aberration frequencies demonstrating, increasing dose dependent increase in all the donors as well as increase in clarity in chromosome dependent sensiviities with increasing aberration data size.

**Table 1.** Chromosomal Specific Aberrations: Pooled Data, 24-donors.

| Dose | Chr1 | Chr2 | Chr4 | Total (X) | Cells (N) |
| --- | --- | --- | --- | --- | --- |
| 0 Gy | 75 | 56 | 44 | 175 | 3452 |
| 2 Gy | 588 | 536 | 395 | 1519 | 2550 |
| 4 Gy | 1569 | 1347 | 976 | 3892 | 2529 |
| Pooled | 2232 | 1939 | 1415 | 5586 | 8531 |

Difference between aberrations from Chr1 and 4 were highly statistically significant at all the dose. Whereas, though aberration yields of Chr2 were also higher than those from Chr4 at all the doses, the difference was marginal at 0 Gy but gained statistical significance at 2 Gy & 4 Gy. Aberration yields between Chr1 & 2 were closer to each other and required larger aberration data to achieve significance. The differences between Chr1 & 2 were marginally significant at 0 Gy and not significant at 2 Gy. However, at 4 Gy and in the combined data, clear statistical significance was observed. The heatmap and scatter plot of individual aberration data also revealed that as we move on from 0 Gy to 2 Gy, 4 Gy and further to pooled data from all the doses, the dominance of Chr1 associated aberration becomes more evident. Except for a few individuals, Chr1 becomes major contributor of the aberrations. Thus, our data suggest that radiation induced aberration susceptibility is highest in Chr1 followed by Chr2 and 4 our population. However, individuals 9 & 20 consistently showed higher aberration frequency for Chr2 than 1 at 2 Gy and 4 Gy. It may be possible to observe higher aberration for Chr2 than 1 in single individual’s data due to statistical reasons associated with limited scoring of aberrations or due to differential radiosusceptibility. When the number of participants is small and the number of aberrations scored is limited, this kind of result is expected. Such data should not be generalized for a population. Consistent with the present study, significantly lower yields of aberration from Chr4 in comparison to Chr1 beyond 2 Gy were recently reported in a G_0_-PCC-FISH study with 3-donors (Yadav et al., 2024). The study could not conclude significant difference between Chr1 & 2, possibly due to small participant numbers. Further, another study reported significantly higher frequency of aberrations from Chr2 than that from Chr1 reporting higher radiosensitivity for the former (Pathak et al., 2009). The study concluded the result based on single donor data but repeated at multiple dose points.

### 3.2. Contribution of different aberration types in total yield: Fragments are predominated aberration types in G_0_-PCC followed by translocations and insertions while Chr1 remains major contributor for all

Overall, 6695 cells were analysed for specific aberrations from 21 donors (21-60Y), yielding 4712 aberrations. The proportions of fragments, translocations, and insertions in these cells were 2,898 (∼61.5%), 1,784 (∼37.8%, 23.8% two-way translocations (TWT), 14% one-way translocations (OWT), and 30 (0.6%), respectively (Figure 3A). Fragments were the predominant type of aberration observed at all the doses, accounting for approximately 71%, 63%, and 61% of the total aberrations in 0 Gy, 2 Gy, and 4 Gy samples, respectively. As the radiation dose increased, higher incidences of translocations and insertions were observed. In the 2 Gy samples, translocations contributed to ∼37% of the total aberrations, which slightly increased to ∼39% in the 4 Gy samples. Notably, the 4 Gy group exhibited a slightly higher proportion of one-way translocations compared to the 2 Gy group.

**Figure 3.**
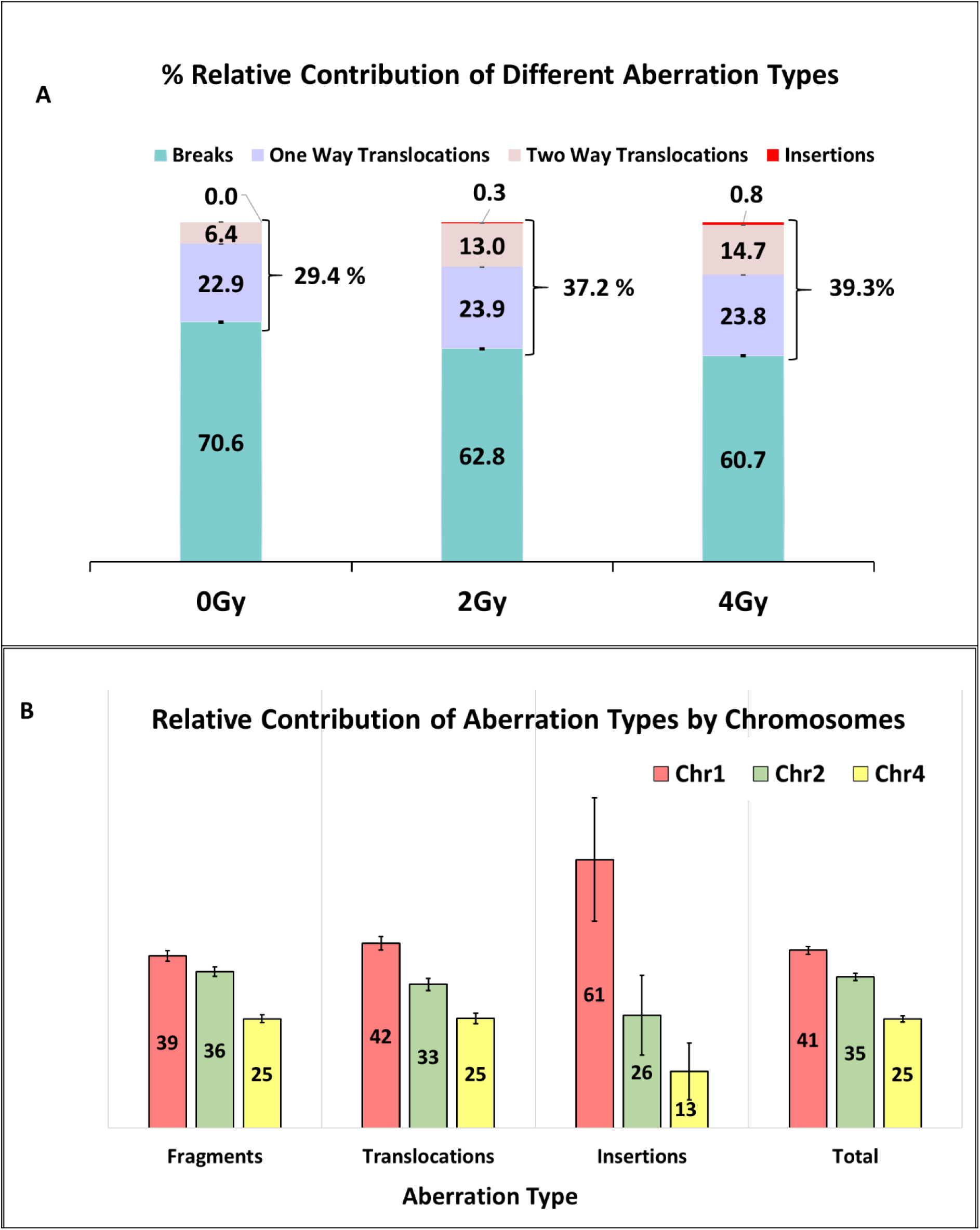
Contribution of different chromosomal aberration types: A. % Contribution of Fragments, Translocations and Insertions to the overall aberration yield at different radiation doses. B. Relative contribution of aberration type chromosome-wise. Fragments can be seen as most predominant type of aberrations followed by translocations & insertions. Chr1 is observed as major contributor for all the aberration types but relatively higher contributions for translocations and insertions.

Insertions, being complex aberrations that involve at least three DSBs to form an interstitial break followed by insertion of the break within a broken chromosome, were observed at relatively low frequencies. No insertions were detected in the 0 Gy samples. In contrast, 4 and 26 insertions were observed in the 2 Gy and 4 Gy samples, respectively, corresponding to only 0.3% and 0.8% of the total aberrations.

Physically, with increase in dose, which inturn proportionately increases number of events per cell, is expected to produce multiple fragments at once leading to higher incidences of misrepairs events such as translocations. Higher doses produce more complex damages, numerous fragments which may also lead to non-reciprocal translocations, one way translocations and in rare cases complex aberrations such as insertions. From physical events point of view, these events are directly proportional target size, which translates to genomic content of each chromosome. Chr1 remained major contributor for all types of aberrations followed by Chr2 & 4. The relative difference was higher for transloactions and insertions compared to the fragments (Figure 3B).

### 3.3. Differential Radiosensitivities among the Chr 1, 2 & 4 and their correlation analysis with density of genes and gene length

To understand the relationship of gene content, gene density and chromosome length on radiosensitivity, we compared the aberration contributions of Chr1, 2 & 4 with their relative size, gene density, gene categories and gene length parameters. The genomic parameters from each chromosome were extracted as described in the method section.

Absolute counts and % contribution of individual chromosomes to the total aberrations are represented in Table 2A. It can be observed from Table 2A that Chr1 is the predominant contributor to aberrations, also predominant in all the gene counts especially, protein-coding genes, tRNA genes and rRNA genes. The chromosome length and gene lengths are also largest in Chr1, except for the non-protein-coding genes and lncRNA transcripts in particular. It is noteworthy that Chr1 being the longest chromosome may be a dominant factor in its contribution to aberrations. Hence, to examine the role of genomic parameters, we normalized the absolute data by the size of the chromosome and compared the aberration density per Million base pairs (Mbp) with the gene densities and gene length densities (per Mbp). The size-normalized data is presented in Table 2B. Even after normalization, aberration density followed the order Chr1 > Chr2 > Chr4, indicating that factors beyond chromosome size may contribute. We generated a correlation matrix to assess the correlation of different genomic parameters with the differential radiosensitivity of the chromosomes (Figure 4; see columns 1 & 2).

**Figure 4.**
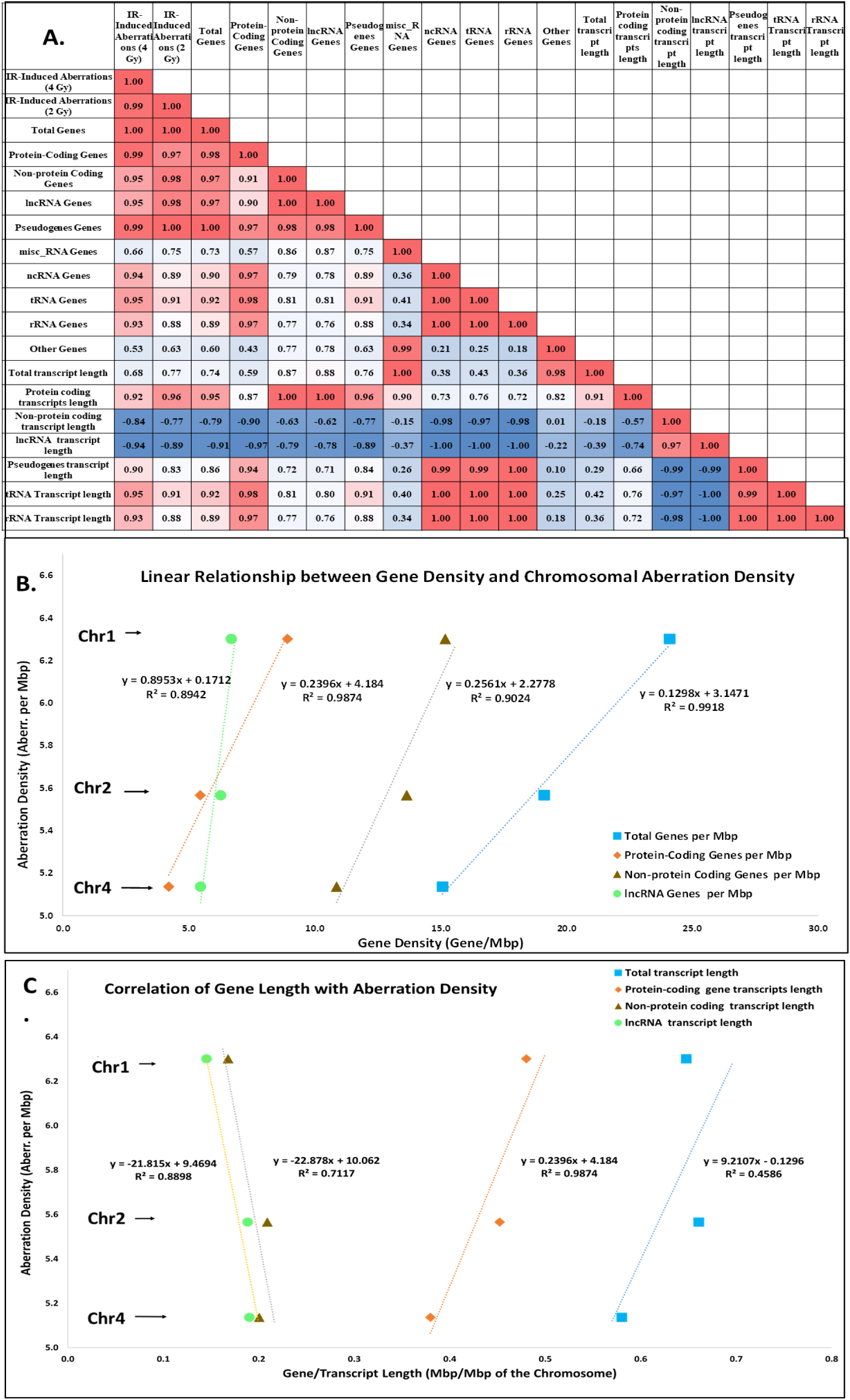
Correlation matrix of IR-induced aberration density with gene density and transcript length across gene categories. This heatmap displays Pearson correlation coefficients (R^2^) between IR-induced chromosomal aberrations (at 4 Gy and 2 Gy) and genomic features, including gene density and gene length, across various gene categories. (B) Regression curve showing linear relationship between gene densities with aberration density as well as with (C) gene length. IR-induced aberration density shows a strong positive correlation with total gene density, protein-coding gene density, and lncRNA gene density. Notably, non-protein coding gene length and lncRNA transcript length exhibits a strong negative correlation with IR-induced aberrations whereas, protein-coding and pseudogene transcript lengths were positively correlated with aberration density.

**Table 2.**
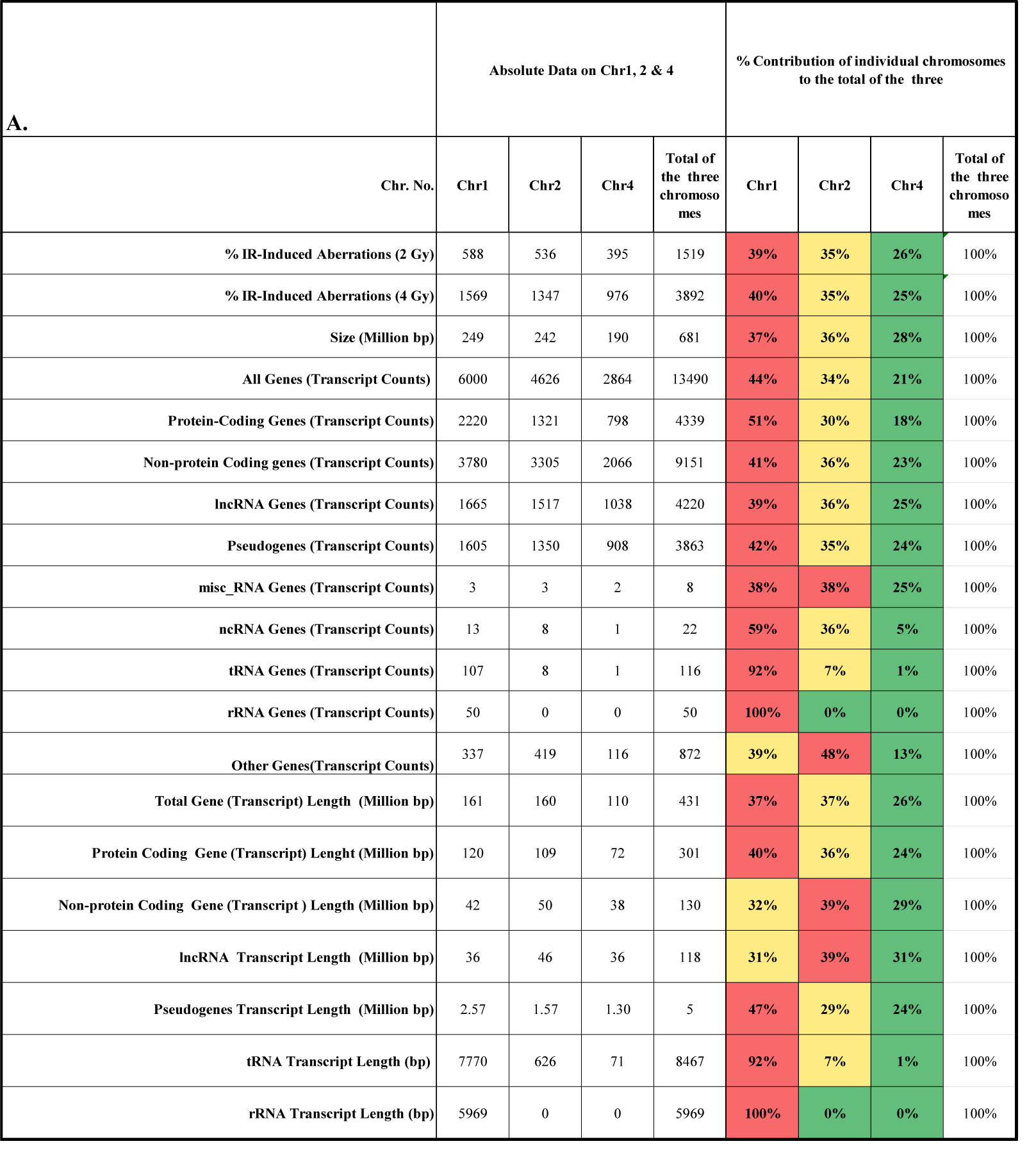
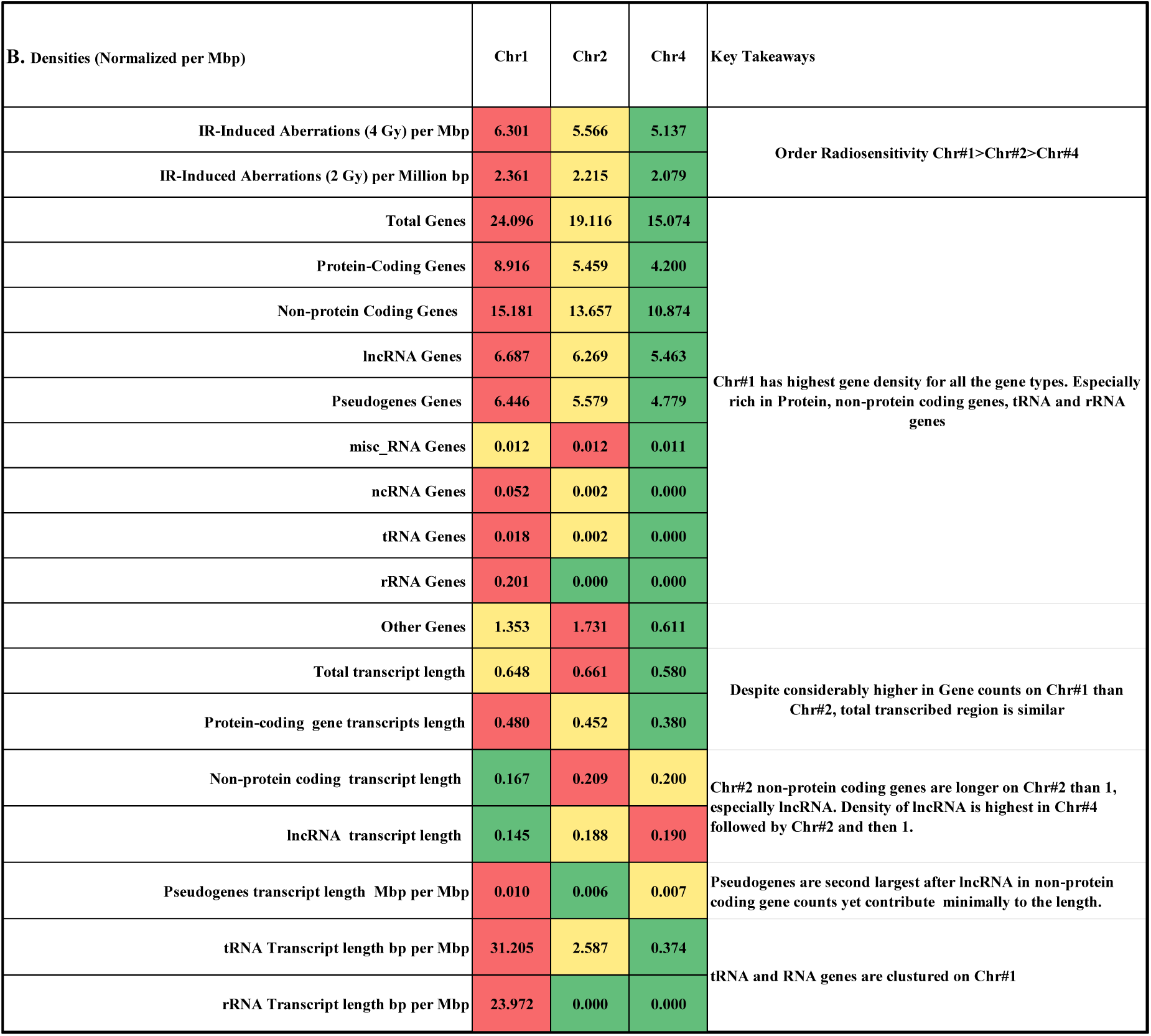
Radiation induced aberrations and genomic features of chromosomes 1, 2, and 4 (A) Absolute values and percentage contributions of chromosomes 1, 2, and 4 to radiation-induced aberrations, gene counts across various gene types, and total transcript lengths. The rightmost columns show the proportion (%) of each chromosome’s contribution to the combined total of all three chromosomes. Gene counts are based on transcript-level annotations, including multiple isoforms and overlapping regions. Corresponding gene-level data are provided in the supplementary data and show similar trends. (B) Densities of aberrations, gene counts, and transcript lengths normalized per million base pairs (Mbp) of respective chromosome.

Chromosome 1 also exhibited the highest gene density across most gene categories and rRNA and tRNA highly expressed genes were clustered on chromosome 1. Regression analysis suggested a positive association between aberration density and total gene density (R² = 0.99, p = 0.058) as well as protein-coding gene density (R² = 0.99, p = 0.072). In contrast, lncRNA transcript length density showed an inverse trend with aberration density (R² = 0.89, p > 0.2). However, due to limitation of analyzing only three chromosomes none of these associations reached statistical significance, and confidence intervals included zero.

These observations suggest that in addition to gene density, non-coding transcript architecture may be associated with chromosome-specific differences in radiosensitivity. Since, current gene counts and lengths are at the transcript level and included total transcript numbers from the chromosome and their respective lengths, we also performed gene level analysis by filtering duplicates which did not change the trend (See supplementary data). Although size is an obvious contributor for susceptibility to a physical clastogen such as radiation, a few reports suggest that chromosomal sensitivity can be disproportional to the size (Barquinero et al., 1998; Knehr et al., 1994, 1996; Luomahaara et al., 1999; Sommer et al., 2005). It is suggested that gene density on a chromosome may be associated with observed differences as these regions are more accessible on chromosome due to their active transcription and hence, higher gene contribution may be related to radiation sensitivity. In an inverse polymerase chain reaction (PCR) based study of selective targets that were rearranged post irradiation in normal human IMR-90 fibroblasts, authors reported that ionizing radiation (IR) induced chromosomal rearrangements occur in transcriptionally active regions of the genome (Forrester & Radford, 2004). It is estimated that ∼80% of the human genome is transcribed and ∼ 98% of the total transcriptional output in humans comes from non-coding DNA (Park et al., 2022; Poliseno et al., 2024). However, lncRNA are known to present with very low expression levels compared to mRNAs (Cabili et al., 2015; Kornienko et al., 2016). Furthermore, in a mice study, lncRNA length and expression levels are reported to be inversely correlated (Li et al., 2013). In a 7 donor study comparison of radiosensitivity of Chr2, 8 and 14 with respect to their size proportions found that Chr14 was unexpectedly contributing to higher aberration in 5 of the 7 analysed donors (Sommer et al., 2005). Whereas, Chr2 was generally less sensitive than expected on the basis of DNA-proportional distribution. We did a supplementary analysis of gene contribution, protein and non-protein coding gene contributions among these three chromosomes and observed that Chr14 despite being smaller, had the highest gene density and smallest lncRNA gene length density with respect to the rest of the two chromosomes (See supplmentary data) Observations from these studies are broadly consistent with the trends observed inour study. Further studies on lncRNA expression, non-coding gene expression in correlation with their length linking to radiosensitivity may enhance current understanding on this topic.

Further, lncRNA expression, non-coding gene expression in correlation with their length linking to radiosensitivity may enhance current understanding on this topic.

### 3.4. Age & Sex Effects: No significant variation with the age (21-60 Y) & sex observed

To understand the influence of age and sex on chromosomal aberrations, combined aberration frequencies arising from the three painted chromosomes were compared among different age groups and the two sexes. We systematically evaluated baseline aberration frequencies and their variation among individuals. Further, to investigate the effect of age and sex on radiosensitivity, 2 Gy and 4 Gy radiation induced aberration frequencies were compared among different age groups (21-60 Y) and the two sexes. A scatter plot of individual aberration frequency and mean aberration frequencies for each age group has been plotted in Figure 5A. Further, the frequency of aberrations across different age intervals is plotted in Figure 5B & C. The sex influence was analyzed after combining all 12 males and 12 females from all age groups as well within age group sex differences were also compared (Figure 6A-E.).

**Figure 5.**
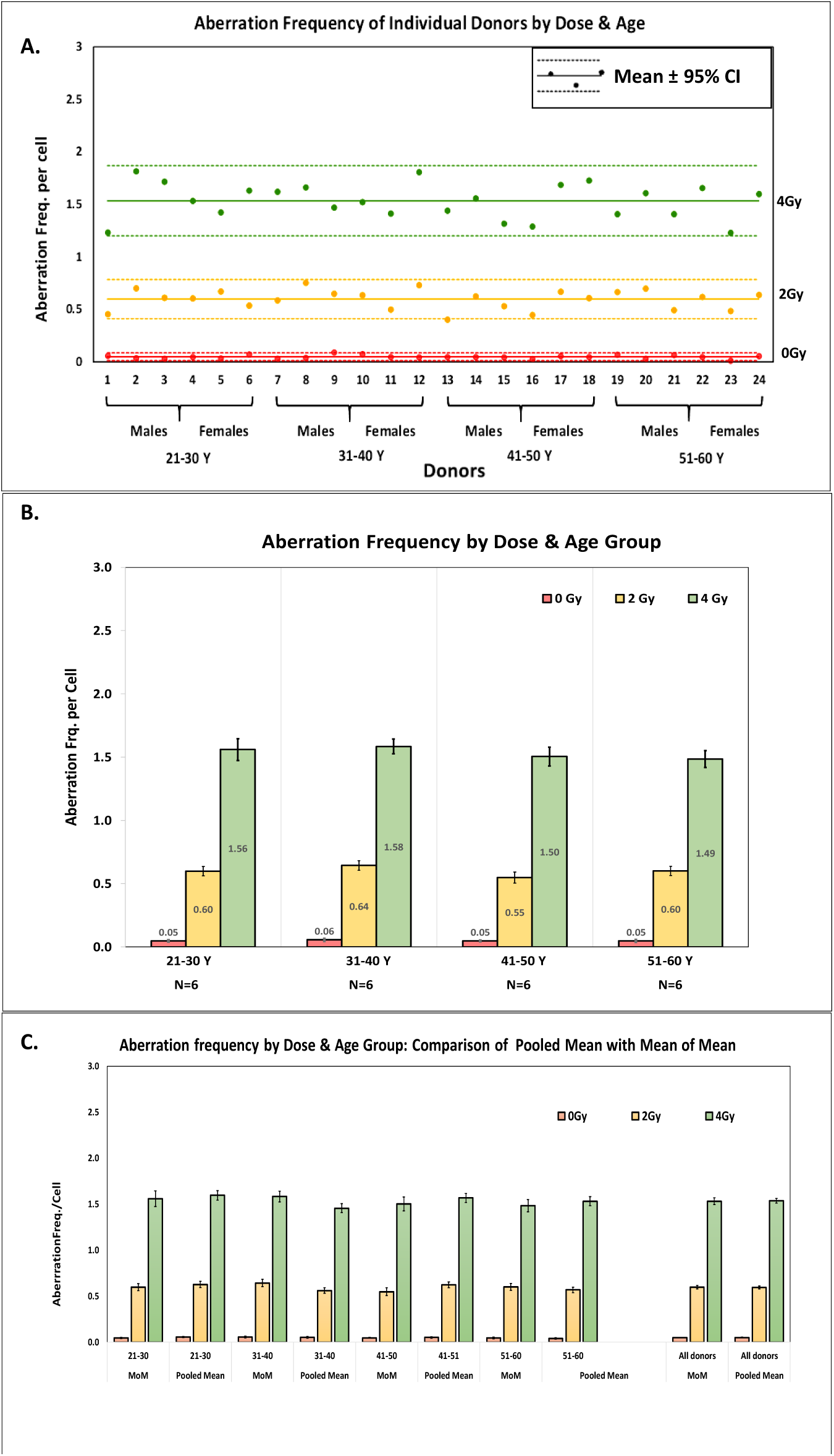
Age and sex variation in baseline and radiation-induced chromosomal aberrations assessed by G₀-PCC-FISH. A) Scatter plot of aberration frequencies at baseline, 2 Gy, and 4 Gy for all 24 donors showed no apparent age-related trend. Solid lines represent mean frequency while dashed line define 95% Confidence Interval. B) B. Aberration frequencies across 4 age intervals presented as Mean ± SEM (6 donors each). No statistically significant differences were observed across age groups at any dose level. C) Comparison of the mean of individual means and the pooled means across age groups and for all 24 donors combined. No significant difference in overall error, indicating consistency between pooled and individual-level analyses.

**Figure 6.**
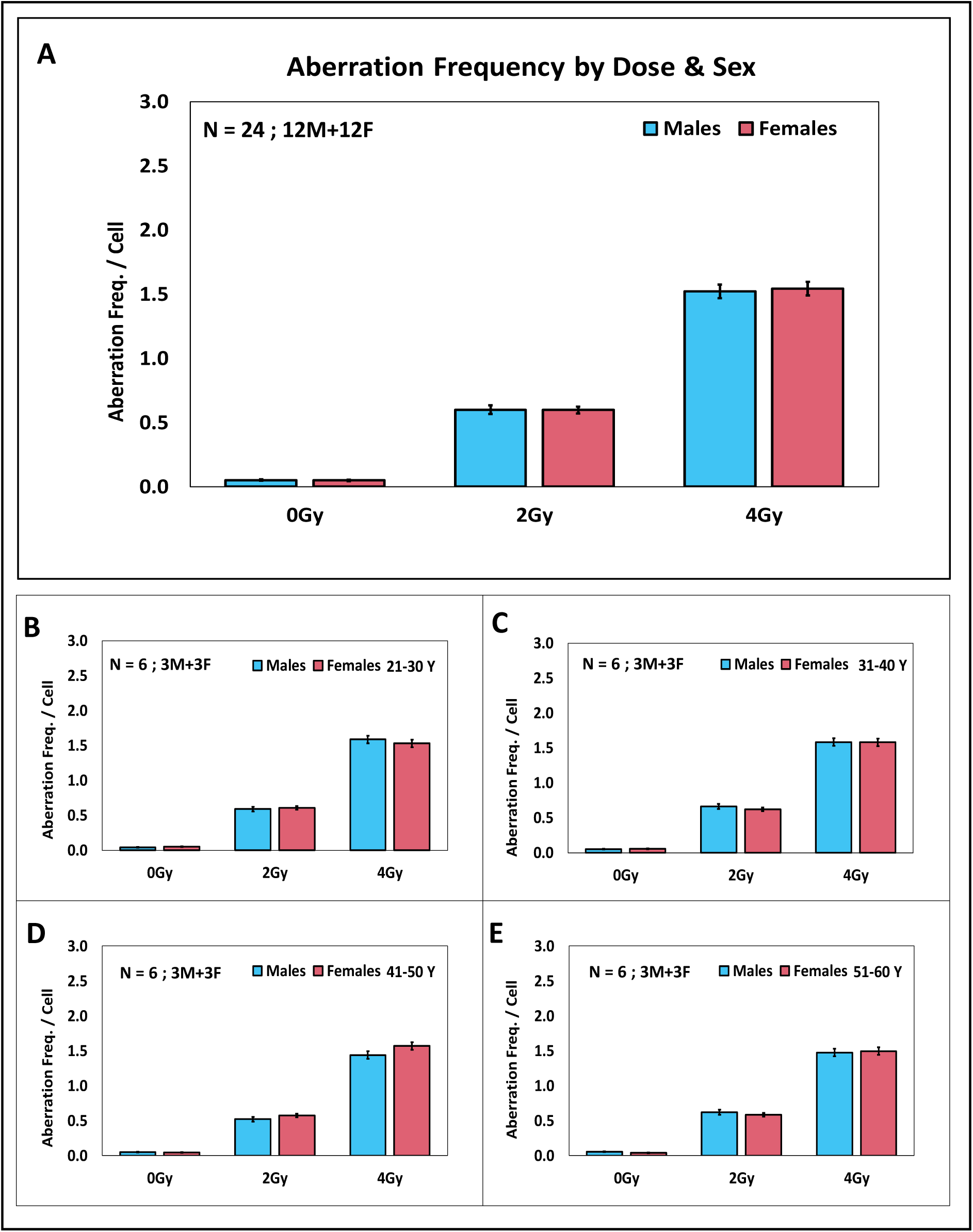
A. Mean frequency of aberrations among males (N = 12) and females (N = 12) showed no significant variation. B-E. Mean frequency of aberration separately plotted for males (N = 3) and females (N = 3) across different age groups, also did not show age dependent difference in radiosensitivity.

#### Baseline Aberrations Variation

The baseline level of aberration in the sample was observed with a mean aberration frequency of 0.049 ± 0.02 per cell, and a median of 0.048 per cell. Further, the frequency of aberrations across different age intervals remained relatively stable, showing no significant variation. For instance, in the 21-30 Y group, the mean aberration frequency was 0.047 ± 0.016 per cell. In the 31-40 Y group, the mean frequency was slightly higher at 0.055 ± 0.024 per cell. For the 41-50 years group, the mean frequency was 0.047 ± 0.010 aberrations per cell, and in the 50-60 years age group, the mean was 0.049 ± 0.02 per cell. The data suggests that age between 21-60 years does not have a significant impact on the aberration frequency.

In terms of sex influence, we also compared the baseline aberration frequencies between males and females. Among the 12 males and 12 females the mean aberration frequency was found to be 0.050 ± 0.019 and 0.048 ± 0.018 per cell respectively. Frequency among females was slightly lower, however, the difference between males and females also was not statistically significant. In conclusion, the data indicates that the aberration frequency is generally low and remains consistent across 21-60 Y age group in both the sexes. This age range is selected with the emphasis on occupational workers, where this method can play a major role for biodosimetry estimations while estimating the minimum detection limit. If the variation is significant, it becomes necessary to have separate response curves for the different age ranges or accept the higher minimum detection limits based on gross variance.

#### Individual Radiation Sensitivity

The mean aberration frequencies induced by 2 Gy and 4 Gy among 24 individuals were 0.60 ± 0.095 and 1.53 ± 0.171, respectively, representing ∼12 - and ∼ 31-fold increases compared to baseline levels. The aberration frequency at 4 Gy was significantly higher than at 2 Gy (p < 0.00001). Both 2 Gy and 4 Gy exposures led to significantly elevated aberration levels in all 24 individuals compared to baseline (p < 0.00001).

No statistically significant differences were observed among the age groups compared to the mean.

Further, sex-based differences in radiation-induced aberration frequencies were assessed among 12 males and 12 females combined from all age groups. At 2 Gy, the mean aberration frequency was 0.60 ± 0.107 in males and 0.60 ± 0.086 in females. At 4 Gy, the frequencies were 1.52 ± 0.172 in males and 1.54 ± 0.176 in females. The values for both sexes were comparable to the overall means, and no statistically significant differences were observed between males and females at either dose level. Earlier studies based on gross count of breaks has indicated a minor increased sensitivity with sex showing marginally higher number of breaks among female samples. The higher yield in gross fragments could be potentially associated with extra X chromosomes in females. However, in the current study, since X-chromosomeis not painted it’s relative contribution is expected to be much lower in the total scored aberration.Additionally, within age group male-female aberration frequencies were also consistent showing no statistical difference. Hence, the pooled age and sex group calibration data can be utilized for dose estimation and it can be ascertained that there is no need to establish age or sex specific response curves.

#### Age and Sex Effects on Individual Chromosomes

Similar to pooled aberration frequencies, no significant influence of age or sex was observed at the level of individual chromosomes. Although some inter-individual variation in chromosome-specific sensitivity was noted, the relative contributions of Chr1, 2, and 4 remained consistent across all age groups and between sexes.

#### Testing individual heterogeneity through comparison of Mean of means with pooled mean

It is essential to assess individual heterogeneity before individual-level data is pooled to make one combined dataset for generating a strong radiation response curve, a common practice for Poisson aberration data. The pooled mean weighs each cell equally, thus, bigger contributors (samples with more cells scored) influence the mean more. In contrast, the mean of individual means treats each individual equally, regardless of how many cells were scored per sample. When we pool data from multiple individuals with significant biological heterogeneity, and then calculate a mean dose-response curve, it essentially smooths out the individual differences, leading to artificially low errors in the mean. Consequently, when estimating dose for a random individual using this pooled curve, it may underestimate the uncertainty of the doses.

A key indicator of between-individual heterogeneity is the comparison of errors: if the pooled standard error of the mean (SEM) is significantly smaller than the SEM of individual means, this suggests meaningful biological variability among individuals.

Towards comparing the individual heterogeneity in our study among 24-donors, we treated per-individual means as a dataset (n = 24) and computed sample mean ± SE. The data set was compared with the pooled mean ± SE (Poisson error) for statistical significance using t-test.

The average mean frequency of aberration across 24 individuals was 0.049 ± 0.004, 0.60 ± 0.019 and 1.52 ± 0.035 for 0 Gy, 2 Gy and 4 Gy respectively. Whereas, corresponding pooled mean of aberrations were 0.050 ± 0.004, 0.60 ± 0.015, 1.56 ± 0.025. The mean aberrations of the individuals were not significantly different from the pooled mean at all the doses. Graphical presentation of Pooled mean ± SE and mean of individual means ± SE are plotted in Figure 5C. It can be observed that the two means as well as the errors are more or less identical. The variability among the individuals is within 95% CI of the pooled means for all the doses. The observation concludes that pooling the data from multiple donors across age groups (21-60 Y) and both the sexes is appropriate and would not affect the accuracy of a calibration curve.

Further, it is to be noted that the age range excludes elderly populations (>65 years) hence all the conclusions are applicable only to the tested age range. Subgroup sizes are modest (n = 6 per age group; n = 3 per sex within age group) hence, statistical power to detect subtle effects may be limited.

## 4. Limitations

It is important to note that correlation/regression analyses presented here are based on only three chromosomes (n = 3), which limits statistical power and robustness. The observed high R² values should therefore be interpreted with caution and considered hypothesis-generating rather than confirmatory. Validation across a larger number of chromosomes may confirm these trends. Further, the present study does not include direct measurements of gene expression, chromatin accessibility, therefore, the proposed links between transcriptional features and radiosensitivity remain correlative. Additionally, overlapping gene annotations were not resolved, which may lead to overestimation of transcript density.

## 5. Conclusions

Present systematic analysis brings out many crucial insights lacking in the literature:

**⁛ Age and Sex independence of aberration (21-60 Y):** Baseline chromosomal aberrations as well as γ-radiation induced aberrations did not show statistically significant variation across age groups within tested range or between sexes indicating the radiation specificity of these aberrations.
**⁛ Chromosome-specific radiosensitivity:** Chr1 exhibited the highest radiosensitivity among 1, 2 & 4 while Chr lowest both in absolute terms and after size normalization.
**⁛ Contribution of different aberration types**; Chromosomal Fragments were the predominant aberration type, followed by translocations and insertions, consistent across all tested chromosomes and doses. Again Chr1 remained major contributor for all types of aberrations.
**⁛ Differential chromosome sensitivities and their correlation with gene density and length:** Correlation analyses suggest a positive association between aberration density and gene density, while an inverse trend is observed with non-protein coding transcript length, particularly lncRNA. These hypothesis are generated based on limited number of chromosomes analysed in the present study.

This study represents first comprehensive analysis of age- and sex-specific chromosomal damage patterns using G_0_-phase PBMCs. From an environmental health perspective, these data provide reference values for assessing radiation-induced chromosomal damage in exposed populations. It enhances emergency preparedness for radiological events by enabling rapid biodosimetry, especially critical when metaphase cells are unavailable as in cases of accidental high-dose or partial exposures.

## 6. Declarations

### 6.1. Ethics approval

Approval from Medical Ethics Committee, Bhabha Atomic Research Centre was obtained before the study approved project number is BHEMC/NP/10/24. All the regulatory guidelines were followed. Informed and written consents were obtained from each of the individual prior to the blood collection. Collection was carried out at the institutional pathology lab by a certified phlebotomist in lithium heparin coated tubes.

### 6.2. Availability of data and materials

The dataset(s) supporting the conclusions of this article is(are) included within the article (and its additional file(s))

### 6.5. Authors’ contributions

U.Y. contributed in conception, design of the work, the acquisition, analysis, interpretation of data, has drafted the work. U.S.M. contributed in majority of data acquisition. N. N. B. contributed in design of the work, analysis, interpretation of the work and substantially revised it. A.K. & B.K.S. substantially revised it. All the authors have approved the submitted version and have agreed both to be personally accountable for the author’s own contributions and to ensure that questions related to the accuracy or integrity of any part of the work, even ones in which the author was not personally involved, are appropriately investigated, resolved and the resolution documented in the literature.

## Acknowledgements

We acknowledge all the donors for their blood sample contribution in the study. Pathology group is being acknowledged for their kind support in blood withdrawal. Mr. Tondlekar P. is being acknowledged for providing extensive technical assistance during the experiments and co-ordination with subjects and pathology teams. In addition, other lab members Mr kapil B Shirsath and Mr. Jagtap S. are for their technical support.

